# Broad gene expression throughout the mouse and marmoset brain after intravenous delivery of engineered AAV capsids

**DOI:** 10.1101/2020.06.16.152975

**Authors:** Nicholas C. Flytzanis, Nick Goeden, David Goertsen, Alexander Cummins, James Pickel, Viviana Gradinaru

## Abstract

Genetic intervention is increasingly explored as a therapeutic option for debilitating disorders of the central nervous system. The safety and efficacy of gene therapies relies upon expressing a transgene in affected cells while minimizing off-target expression. To achieve organ/cell-type specific targeting after intravenous delivery of viral vectors, we employed a Cre-transgenic-based screening platform for fast and efficient capsid selection, paired with sequential engineering of multiple surface-exposed loops. We identified capsid variants that are enriched in the brain and detargeted from the liver in mice. The improved enrichment in the brain extends to non-human primates, enabling robust, non-invasive gene delivery to the marmoset brain following IV administration. Importantly, the capsids identified display non-overlapping cell-type tropisms within the brain, with one exhibiting high specificity to neurons. The ability to cross the blood–brain barrier with cell-type specificity in rodents and non-human primates enables new avenues for basic research and potential therapeutic interventions unattainable with naturally occurring serotypes.

## INTRODUCTION

The wide array of debilitating disorders affecting the central nervous system (CNS) make it a primary target for the application of novel therapeutic modalities, as evidenced by the rapid development and implementation of gene therapies targeting the brain. Adeno-associated viruses (AAVs) are increasingly utilized as delivery vehicles due to their strong clinical safety record, low pathogenicity, and stable, non-integrating expression *in vivo*^1^. AAVs were first approved for gene therapy in humans in 2012 to treat lipoprotein lipase deficiency^2^, and have more recently been approved to treat spinal muscular atrophy^3^, retinal dystrophy and hemophilia^4,5^. Many more advanced-stage clinical trials using AAVs are underway^6–11^. Currently, most AAV-based gene therapies rely on naturally occurring serotypes with highly overlapping tropisms^12^, limiting the applicability, efficacy, and safety of novel gene therapies. At the same time, the inability to broadly and efficiently target many relevant therapeutic populations for genetic delivery has hindered novel exploratory and therapeutic efforts in organs that have traditionally been refractory to AAV delivery. These constraints, which are especially prevalent for the CNS, motivated us to enhance AAV efficiency and specificity through directed evolution.

Naturally occurring AAVs have evolved to infect cells broadly^13^, a desirable characteristic for the survival of the virus but undesirable for targeting specific cell types. This limitation has typically been addressed by injecting viral vectors directly into the area of interest in the CNS^14^, either through intracranial (IC) or intrathecal (IT) injection. This approach greatly limits the efficacy of therapeutic applications since direct injections are highly invasive, offer limited coverage, and require considerable surgical expertise^14–16^. Systemic administration into the bloodstream is an appealing solution to the limitations of direct injections, but the naturally occurring AAV serotypes tend to target non-CNS tissues, such as the liver, at high levels^12,13^, and have severely limited transduction efficiency in the brain due to the stringency of the blood–brain barrier. Previously, we harnessed the power and specificity of Cre transgenic mice to apply increased selective pressure to viral engineering, leading to the variants AAV-PHP.B and AAV-PHP.eB, which cross the blood–brain barrier following intravenous (IV) administration and broadly transduce cells throughout the CNS^17,18^. These variants transformed the way we study the CNS in rodents, opening up completely novel approaches for measuring and affecting brain activity, both in exploratory and translational contexts^19–22^. However, these variants also transduce cells in the liver and other off-target organs. The liver is an immunologically active organ, with large populations of phagocytic cells that play a critical role in immune activation^23,24^, making liver detargeting particularly important for avoiding strong, systemic immune responses^25–27^.

To achieve specificity, it is imperative that one applies both positive and negative selective pressure to engineering capsid libraries. By utilizing next-generation sequencing (NGS) of synthetic libraries with built-in controls and by screening across multiple Cre transgenic lines for both positive and negative features using the M-CREATE method^28^, we selected a panel of novel variants that are both highly enriched in the CNS and detargeted from varying peripheral organs. Most notably, AAV.CAP-B10 was found to be highly specific for neurons in the CNS, significantly detargeted from all peripheral organs assayed, and nearly completely detargeted from the liver in mice. Importantly, while previously engineered variants for CNS transduction did not translate to non-human primates (NHPs)^29,30^, here we show robust DNA, RNA, and protein expression of newly engineered variants in the adult marmoset CNS following IV administration, with AAV.CAP-B10 and AAV.CAP-B22 resulting in ~19-fold and ~40-fold increases in transgene levels in the brain compared with AAV9, respectively.

## RESULTS

### Engineering AAV capsids at the 3-fold point of symmetry

The most commonly manipulated position within the AAV capsid is the surface-exposed loop containing amino acid (AA) 588 because it is the site of heparan sulfate binding in AAV2^31^ and is amenable to peptide display^32,33^. The only known receptors for AAV9 are N-linked terminal galactose^34^ and the AAV receptor (AAVR)^35,36^, but the possibility of co-receptors is still unexplored. Binding interactions with cell-surface receptors occur near the 3-fold axis of symmetry of the AAV9 viral capsid (Fig. 1a) where the AA455 loop is the furthest protruding (Fig. 1b)^37^. This loop has previously been implicated in neutralizing antibody binding^38^. Having already engineered AAV-PHP.B and AAV-PHP.eB at the 588 loop for enhanced CNS transduction^17,18^, and with the further goal of detargeting viral transduction from peripheral organs, we theorized that introducing diversity into the AA455 loop would enhance the interaction with existing mutations of the 588 loop (Fig. 1c) and refine transduction. Thus, we engineered and performed two rounds of selection with a 7-amino-acid substitution library of the 455 loop, between AA452-458 (Fig. 1a-c), in AAV-PHP.eB.

**Figure 1.**
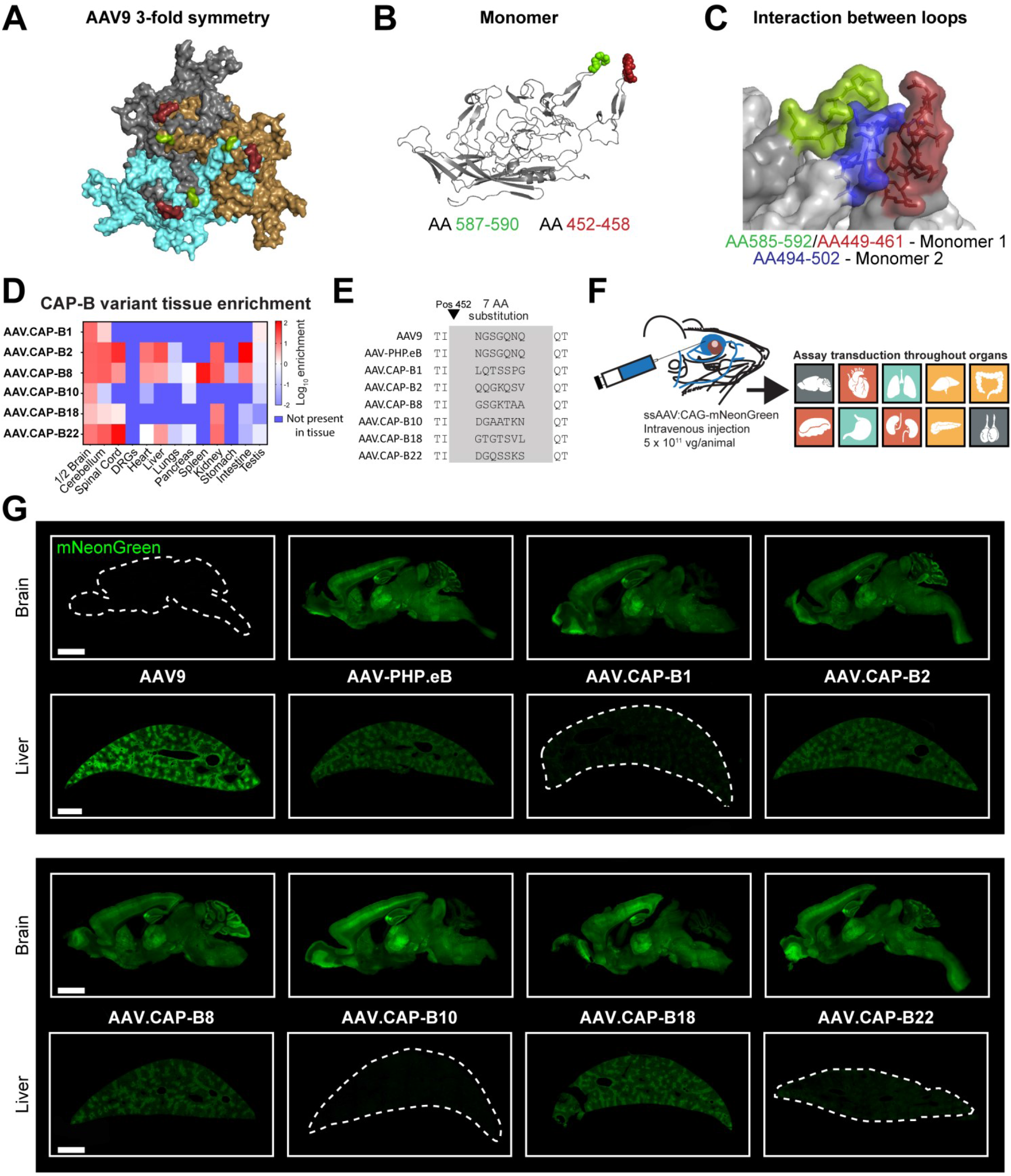
Capsid engineering locations and CAP-B library characterization in mice. (**a**) Surface model of the three monomers comprising the 3-fold symmetry of AAV9, illustrating the location of the 7-amino-acid (AA) substitution introduced in this library (red) and the 2-AA substitution and 7-AA insertion introduced by PHP.eB previously (green). (**b**) Position of the two neighboring surface loops within a single AAV9 monomer. (**c**) Spike created by the 588 and 455 loops of one AAV9 monomer interacting with the 495 loop of a second monomer. (**d**) Heat map plotting the enrichment scores of a subset of the top performers after two rounds of selection *in vivo* demonstrates enrichment and specificity for the CNS. (**e**) Sequences for the 7-AA substitutions present in each of the variants from (**d**) show strong divergence from the WT AAV9 sequence. See also Figure S1. (**f**) mNeonGreen was packaged in each variant under control of the ubiquitous CAG promoter and intravenously administered to mice at a dose of 5 × 10^11^ vg/animal. Transduction was assayed by mNeonGreen fluorescence throughout organs. (**g**) Expression from ssAAV9, ssAAV-PHP.eB, ssAAV.CAP-B1, ssAAV.CAP-B2, ssAAV.CAP-B8, ssAAV.CAP-B10, ssAAV.CAP-B18, and ssAAV.CAP-B22 packaging CAG-mNeonGreen as assessed after two weeks. Direct comparison of the expression profiles in brain and liver of the top-performing variants shows a strong correlation between validated tropisms and those predicted by the NGS data. Scale bars are 2 mm.

To screen for AAVs specific for the CNS following systemic administration, we applied the M-CREATE method developed by Kumar and colleagues^28^, which enables simultaneous positive and negative selection of large libraries of viral variants *in vivo*. With M-CREATE, a library of AAV capsids is generated by mutating a specific location and viruses are produced in HEK293 cells that package a replication-incompetent version of their own genome with a polyadenylation sequence flanked by Cre/Lox sites. The viral library is then injected into transgenic animals expressing Cre in a specific population of cells. Variants that successfully transduce Cre+ cells have their genome flipped, and the sequences of only those variants are recovered. Diverse capsid sequences were identified through next generation sequencing (NGS). Variants in viral libraries can be selected for specificity by using multiple transgenic mouse lines expressing Cre in different populations — Tek-Cre for endothelial cells throughout the body, hSyn1-Cre for neurons of the CNS and peripheral nervous system (PNS), and GFAP-Cre for astrocytes — and in wild-type (WT) mice as a control. The results were run through a bioinformatics pipeline, mining the datasets for sequences that were positively enriched in tissues of interest (the CNS in this case) and negatively enriched in others. We then ranked sequences based on these criteria and narrowed the library down to only the top 10% of variants.

### Iterative capsid engineering refines expression patterns in mice

The first round of selection yielded over 82,000 unique variants that were found to be enriched in the CNS with varying patterns of detargeting from peripheral organs. We used this dataset to synthesize a library for the second round of selection containing these codon sequences, and a duplicate, codon-modified version, as an internal control. The second round of selection narrowed down the top-performing variants by two orders of magnitude and differentiated a subset of AAs that contributed to high CNS enrichment while detargeting from the liver (Fig. S1). Using the NGS data, we isolated a small subset of sequences with high enrichment in the CNS and variable enrichment throughout the periphery (Fig. 1d, e) for testing *in vivo*. We tested this subset individually in WT mice by intravenously injecting 5 × 10^11^ viral genomes packaging CAG-mNeonGreen, allowing two weeks for expression, and then assaying transduction throughout all organs of the body by fluorescence (Fig. 1f). The resulting expression in the brain and liver (Fig. 1g) correlated very closely with the NGS enrichments; variant AAV.CAP-B10 stood out by exhibiting higher fluorescence in the brain than AAV-PHP.eB as well as negligible liver transduction. Additionally, AAV.CAP-B22 was notable for having the strongest brain transduction of any of the variants tested *in vivo*.

### AAV.CAP-B10 exhibits CNS-specific transduction in mice

To fully characterize the performance of AAV.CAP-B10 in comparison with AAV9 and AAV-PHP.eB, we packaged a nuclear-localized CAG-EGFP, injected it IV into mice at a dose of 1 × 10^11^ viral genomes, and allowed three weeks for expression. Detailed cell counts of overall transduction throughout the body showed that AAV.CAP-B10 is highly specific to the CNS. We found a non-significant increase in the average number of cells transduced in the brain, as well as an increased average level of expression per cell, compared with AAV-PHP.eB; both AAV.CAP-B10 and AAV-PHP.eB had much higher transduction of the brain than AAV9 (Fig. 2a). In the spinal cord, AAV.CAP-B10 transduced approximately 40% fewer cells than AAV-PHP.eB, but approximately 16-fold more than AAV9 (Fig. S2). Conversely, AAV.CAP-B10 transduction was very significantly reduced in the liver compared with both AAV-PHP.eB (~50-fold) and AAV9 (>100-fold). In the liver, AAV.CAP-B10 expression, but not AAV-PHP.eB expression, was significantly dimmer per cell than AAV9 (~10-fold) (Fig. 2b). This trend was maintained in the other peripheral organs, with AAV.CAP-B10 transducing significantly fewer cells than AAV9 (Fig. S2).

**Figure 2.**
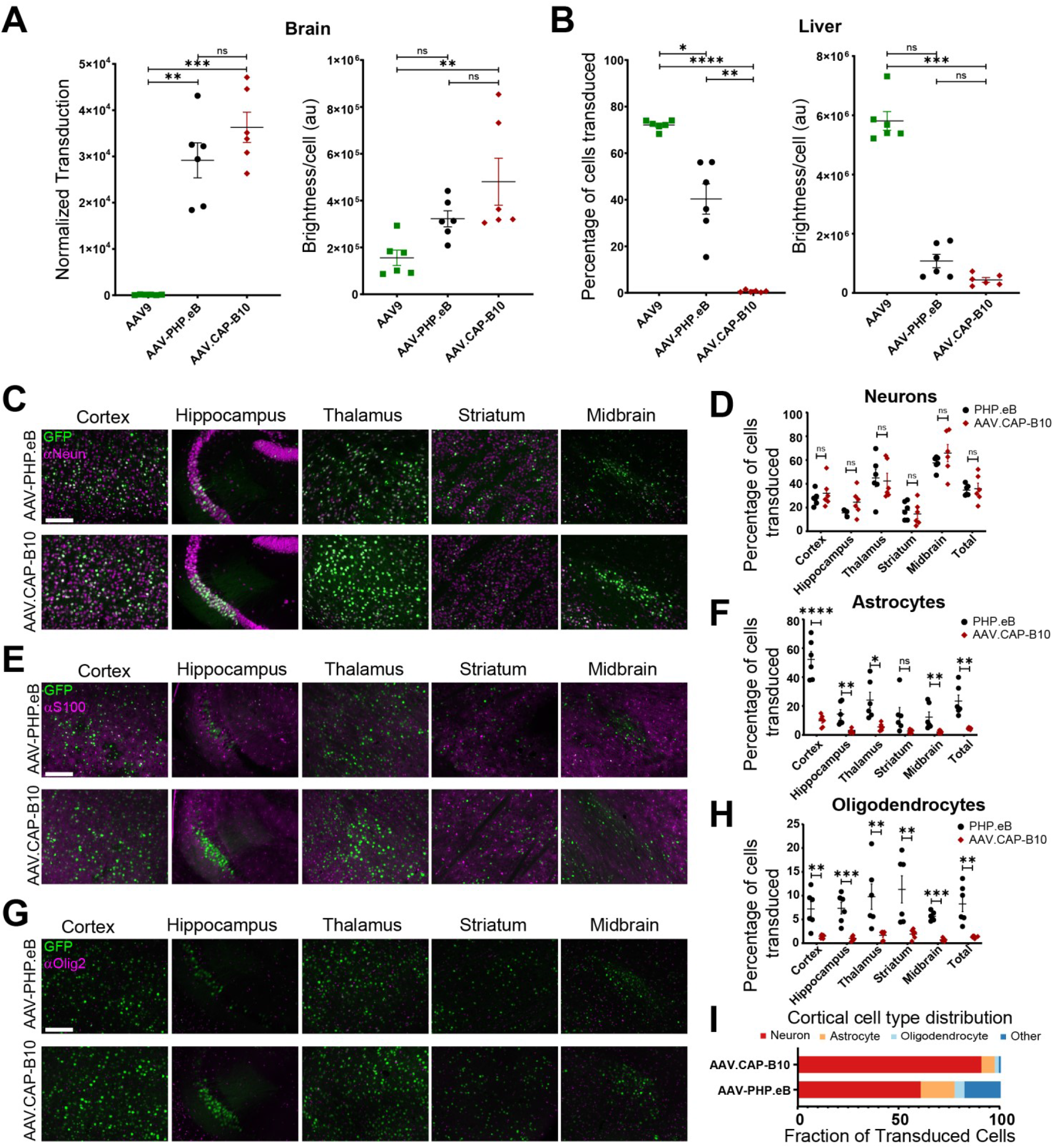
AAV.CAP-B10 tropism in mice is strongly biased towards the brain, with significant liver detargeting. ssAAV9, ssAAV-PHP.eB, and ssAAV.CAP-B10 packaging a nuclear localized GFP under control of the CAG promoter were intravenously injected into male adult mice at 1 × 10^11^ vg/mouse. GFP fluorescence was assessed after three weeks of expression. (**a**) Quantification of the total number of cells transduced in the brain and the average brightness per cell shows a progressive increase in transduction efficiency in the brain. (**b**) Quantification of the total percentage of cells transduced in the liver and average brightness per cell shows a progressive decrease in transduction efficiency. See also Figure S2. (**a**, **b**) n = 6 mice per group, mean ± SE, Brown-Forsythe and Welch ANOVA tests for transduction and Kruskal-Wallis test for brightness. (**c-h**) Within the brain, AAV.CAP-B10 is strongly biased towards neurons. n = 6 mice per group, mean ± SE, multiple t tests. (**c, d**) Expression and percentage of neurons transduced. (**e, f**) Expression and percentage of astrocytes transduced. (**g, h**) Expression and percentage of oligodendrocytes transduced. (**i**) Cortical cell-type bias of AAV.CAP-B10 and AAV-PHP.eB. EGFP expression in four areas of the mouse cortex was co-localized with cell-type-specific staining for neurons, astrocytes, and oligodendrocytes. The contribution of those cell-types to overall EGFP expression was measured and indicates a heavy shift toward neuronal specificity for AAV.CAP-B10 compared with AAV-PHP.eB. See also Figure S3 and Figure S4. Scale bars are 200 μm.

### AAV.CAP-B10 transduction in the CNS is highly specific to neurons

To further characterize CNS transduction by AAV.CAP-B10 compared with AAV-PHP.eB, we co-stained for neurons, astrocytes, oligodendrocytes, and Purkinje cells and quantified the transduction efficiency of each capsid for each cell type across various brain regions (Fig. 2c-i, Fig. S3). Whereas neurons were transduced with similar efficiencies by AAV.CAP-B10 and AAV-PHP.eB across brain regions (Fig. 2c, d), astrocytes and oligodendrocytes were transduced at roughly 4-to 5-fold lower levels by AAV.CAP-B10 compared with AAV-PHP.eB (Fig. 2e-g). This indicates that the AAV.CAP-B10 mutation confers a bias for neurons over other cell types, an interesting deviation from AAV9, which mostly targets astrocytes in the brain^39,40^. A noteworthy indication from the NGS data for AAV.CAP-B10 (Fig. 1d) was this variant’s negative enrichment in the cerebellum, which was also evidenced by a decrease in the overall cerebellar fluorescence in the initial characterization (Fig. 1g). When studying the cerebellar expression of AAV.CAP-B10 compared with that of AAV-PHP.eB in more detail, we did indeed find a significant, roughly 4-fold, decrease in the transduction of Purkinje cells (Fig. S4).

### AAV.CAP-B variants maintain robust neurotropic transduction and cell-type bias in marmosets

Of primary concern for the therapeutic applicability of capsid variants engineered in rodents is how well their transduction profiles translate to NHPs. Therefore, we sought to characterize the marmoset CNS transduction of a subset of the variants validated in mice, together with AAV9 and AAV-PHP.eB as controls. We produced a pool of eight viruses – AAV9, AAV-PHP.eB, AAV.CAP-B1, AAV.CAP-B2, AAV.CAP-B8, AAV.CAP-B10, AAV.CAP-B18, and AAV.CAP-B22 – each packaging an HA-tagged frataxin (FXN) with a unique molecular barcode under control of the ubiquitous CAG promoter. We opted to use FXN because it is an endogenous protein expressed throughout the body; previous efforts to characterize NHP transduction of naturally occurring and engineered serotypes have resulted in deleterious effects on the tissue, potentially due to the packaging of exogenous transgenes such as GFP^25,41,42^. Each packaged FXN contained a separate 12-base RNA barcode to differentiate the contribution of each virus from the rest after sequencing. The eight viruses were pooled at equal ratios and injected IV into two adult marmosets at doses of 1.2 × 10^14^ (MPV1) and 7 × 10^13^(MPV2) vg/kg (Fig. 3a, Table 1). Following six weeks of expression, throughout which no adverse health effects were observed, the brains were recovered, and coronal sections taken for RNA sequencing and immunohistochemistry. Livers were also subjected to RNA sequencing.

**Figure 3.**
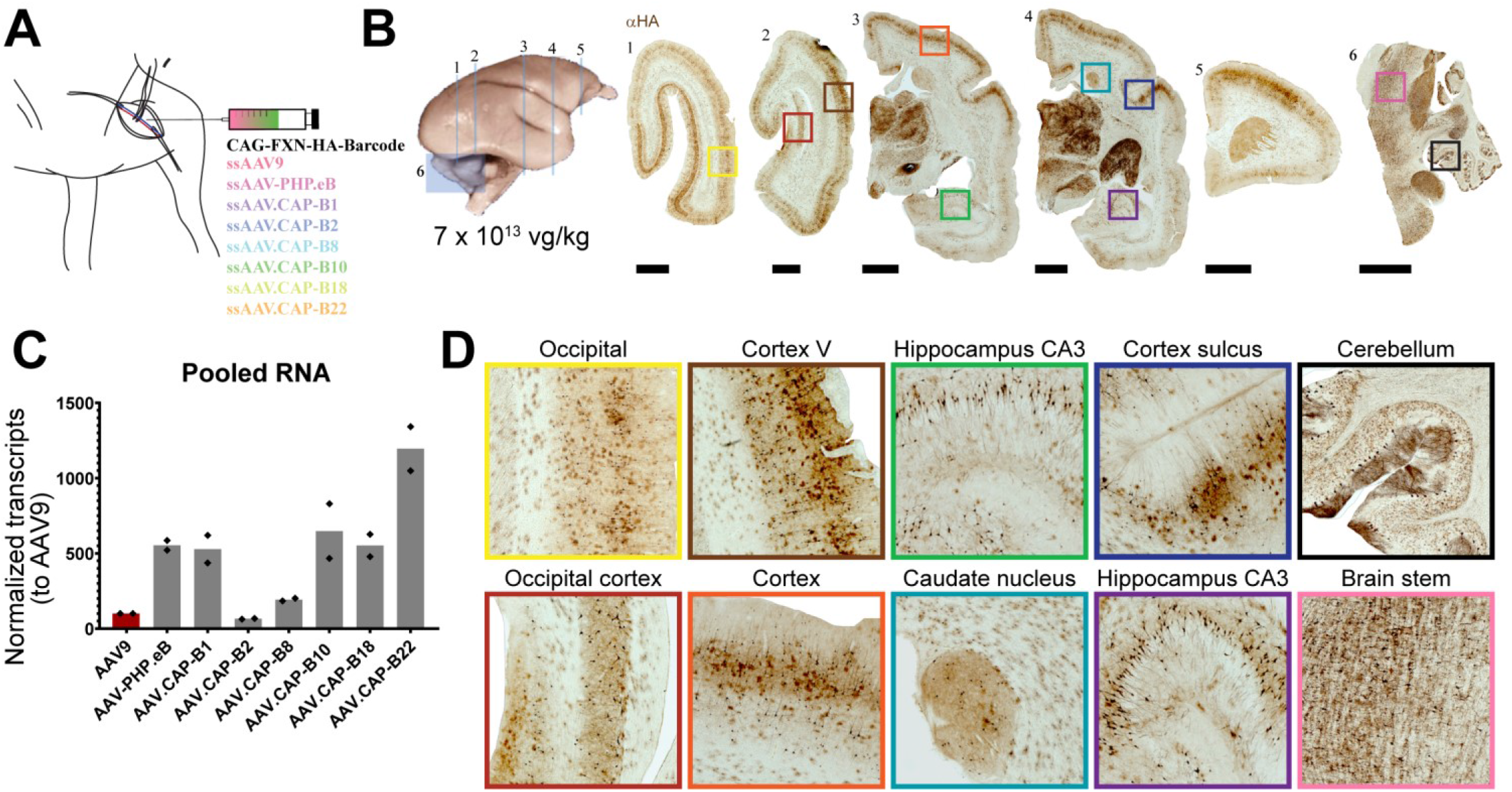
Pooled expression in marmosets. (**a**) AAV.CAP-B1, AAV.CAP-B2, AAV.CAP-B8, AAV.CAP-B10, AAV.CAP-B18, and AAV.CAP-B22, along with AAV9 and AAV-PHP.eB as controls, packaging a human frataxin (FXN) fused to an HA tag under control of the ubiquitous CAG promoter, were pooled and intravenously injected into two adult marmosets at doses of 1.2 × 10^14^ (MPV1) and 7 × 10^13^ (MPV2) vg/kg in total. (**b**) Six sections distributed throughout the brain of one marmoset (MPV2) show robust expression after immunostaining. (**c**) NGS quantification within brain tissue of the unique RNA barcode associated with each virus shows a dramatic increase for several variants, including a >12-fold increase in RNA levels of AAV.CAP-B22 and a >5-fold increase for AAV.CAP-B10 compared with AAV9. See also Figure S5. (**d**) Magnified frames from (**b**) for a variety of cortical and sub-cortical regions. n = 2 marmosets. Scale bars are 2 mm.

**Table 1.** Systemic marmoset injections of a CNS-targeting engineered viral pool. MPV stands for Marmoset Pool Virus.

| **MARMOSET**<br>**ID#** | **AGE**<br>**(YEARS)** | **SEX** | **WEIGHT AT**<br>**INJECTION (KG)** | **DOSE (VG/KG)** | **EXPRESSION**<br>**(DAYS)** |
| --- | --- | --- | --- | --- | --- |
| MPV1 | 7.6 | male | 0.386 | $1.2 \times 10^{14}$ | 42 |
| MPV2 | 11.5 | male | 0.375 | $7 \times 10^{13}$ | 42 |

Robust and broad expression was achieved in the adult marmoset brain, with strong expression of the viral pool in cortical, subcortical, and cerebellar layers throughout (Fig. 3b, d). Following RNA extraction and NGS from multiple coronal sections per animal distributed throughout the brain, as well as sections from the liver, we quantified the relative expression levels of each of the barcoded viruses. AAV.CAP-B10 recapitulated the trends observed in mice (albeit at lower levels), with a ~6-fold increase in RNA in the brain and a ~5-fold decrease in RNA in the liver compared with AAV9 (Fig. 3c, Fig. S5a). AAV.CAP-B22 exhibited the largest increase in RNA levels in the brain, with a greater than 12-fold increase in the brain compared with AAV9, but relatively similar levels in the liver (Fig. 3c, Fig. S5a). Based on their unique transcription profiles in the pooled screen, AAV.CAP-B10 and AAV.CAP-B22 were selected for individual characterization of their CNS transduction in adult marmosets compared with AAV9.

Individual adult marmosets were injected IV with either AAV9, AAV.CAP-B10, or AAV.CAP-B22 at a dose of 7 × 10^13^ vg/kg (Table 2). Following thirty-four days of expression, the brains were recovered, and coronal sections were taken for DNA sequencing, RNA sequencing, and immunohistochemistry. To characterize CNS transduction of the three variants, sections were stained for the presence of the HA-tag. In stained sections, both AAV.CAP-B10 and AAV.CAP-B22 show a marked increase in transduction efficiency within cortical and sub-cortical regions compared with AAV9 (Fig. 4a and b).

**Table 2.** Systemic marmoset injections of CNS-targeting engineered viral variants. MPS stands for Marmoset Single Variant.

| **MARMOSET**<br>**ID#** | **AGE**<br>**(YEARS)** | **SEX** | **WEIGHT AT**<br>**INJECTION (KG)** | **VARIANT**<br>**INJECTED** | **DOSE**<br>**(VG/KG)** | **EXPRESSION**<br>**(DAYS)** |
| --- | --- | --- | --- | --- | --- | --- |
| MSV 1 | 11.5 | female | 0.300 | AAV9 | $7 \times 10^{13}$ | 34 |
| MSV 2 | 6.9 | female | 0.450 | AAV.CAP-B10 | $7 \times 10^{13}$ | 34 |
| MSV 3 | 12.1 | male | 0.360 | AAV.CAP-B10 | $7 \times 10^{13}$ | 34 |
| MSV 4 | 14.0 | male | 0.400 | AAV.CAP-B22 | $7 \times 10^{13}$ | 34 |

**Figure 4.**
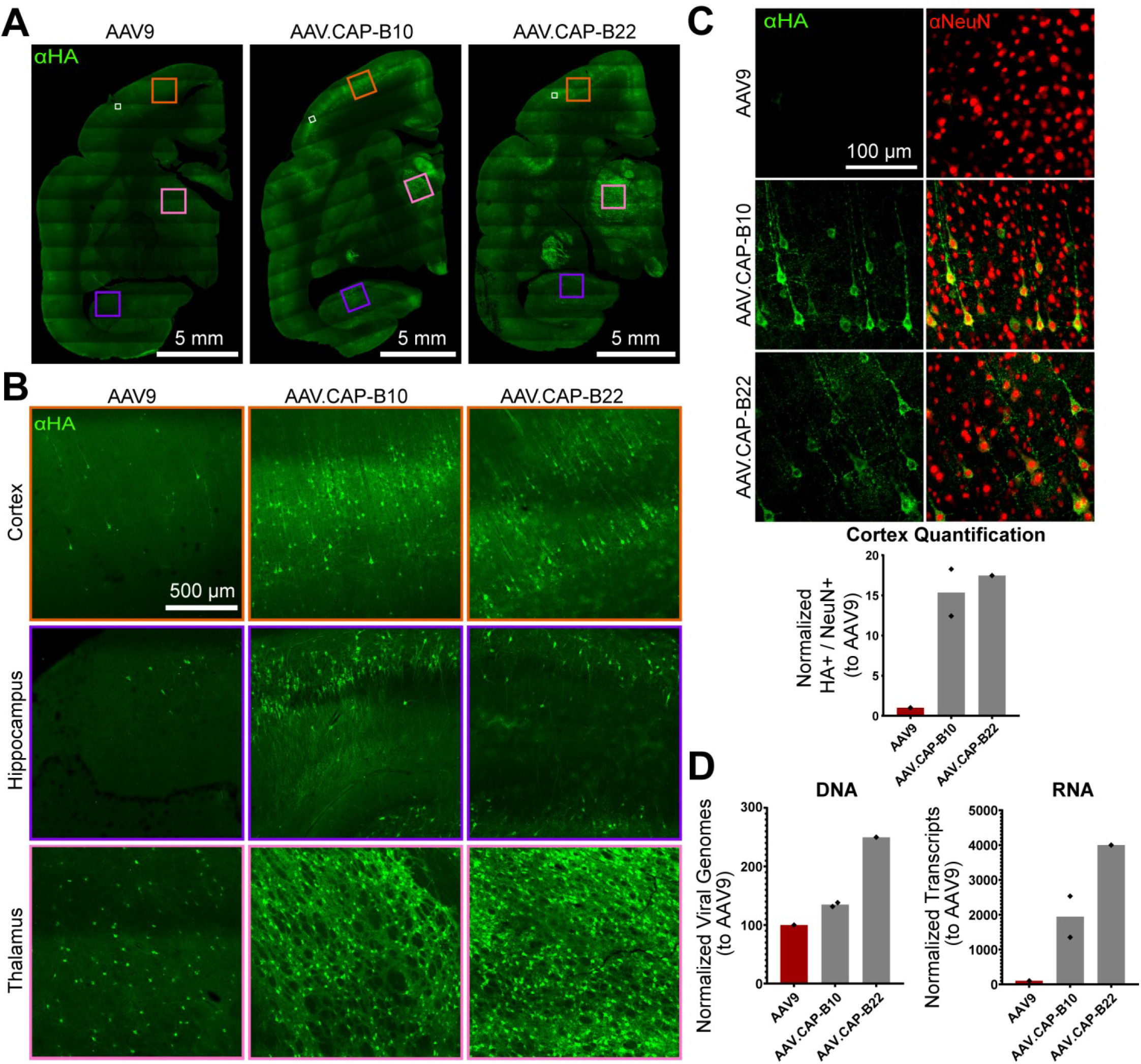
Single variant expression of AAV9, AAV.CAP-B10, and AAV.CAP-B22 in marmosets. Human frataxin (FXN) fused to an HA tag was packaged in each variant under control of the ubiquitous CAG promoter. Marmosets were injected at a dose of 7 × 10^13^ vg/kg. See also Figure S6. (**a**) Coronal sections were taken from a similar plane across animals and AAV9, AAV.CAP-B10, and AAV.CAP-B22 transduction was compared by immunostaining the sections for the HA tag on the FXN transgene. AAV.CAP-B10 and AAV.CAP-B22 show increased transduction efficiency compared with AAV9. (**b**) Magnified frames from cortex, hippocampus, and thalamus show increased transduction in each examined region for both engineered variants in comparison to AAV9. (**c**) Neuronal transduction was manually quantified in layer 3 of the marmoset cortex. AAV.CAP-B10 and AAV.CAP-B22 show a ~15-fold and a ~17-fold increase in HA-positive neurons over AAV9, respectively. (**d**) Viral genomes and RNA transcripts were measured in brain tissue for each capsid variant: AAV.CAP-B10 and AAV.CAP-B22 both show higher transduction in the brain than AAV9. See also Figure S5. Total values for viral genomes were normalized by tissue mass, while RNA transcripts were normalized by the total number of transcripts. Scale bar lengths are indicated on the associated images.

The neuronal specificity of AAV.CAP-B10 in mice was recapitulated in marmoset brains, where HA-positive cells were highly correlated with NeuN staining across brain regions. By contrast, AAV.CAP-B22 had broad cell-type transduction in the marmoset brain, transducing many more astrocytes than either AAV.CAP-B10 or AAV9 (Fig. S6). Both engineered variants showed a substantial increase in the number of neurons transduced in layers 3 and 5 of the marmoset cortex relative to AAV9, with AAV.CAP-B10 and AAV.CAP-B22 displaying ~15-fold and ~17-fold increases in HA-positive neurons, respectively (Fig. 4c). Following DNA and RNA extraction in the brain and liver, normalized transduction and expression measurements corroborate the protein expression data (Fig. 4d, Fig. S5b): AAV.CAP-B10 and AAV.CAP-B22 displayed ~19-fold and ~40-fold increases in transgene expression in the brain, respectively, relative to AAV9.

## DISCUSSION

The power of directed evolution and AAV engineering to confer novel tropisms and cell-type specificity has broadened potential research applications and enabled new therapeutic approaches in the CNS^43–48^. Our results here show that introducing diversity at multiple locations on the capsid surface into native or previously engineered variants can result in useful features, such as increased transduction efficiency and tissue or cell-type specificity in wild-type (WT) mice and marmosets. This finding has broad implications for the field of AAV engineering, and the variants discovered using the M-CREATE method^17,18,28^ have the potential to provide cell-type-specific gene delivery in WT animals.

Novel variants from this library achieved overall similar levels of transduction throughout the brain as the previously engineered parent AAV (AAV-PHP.eB), but with striking deviations in cell-type specificity and targeting or, importantly, de-targeting of other organs. Across the two engineering efforts, iterative modifications made to adjacent loops first conferred a strong new phenotype – the ability to cross the blood–brain barrier efficiently – and then refined the phenotype away from other organs or toward specific cell types. Most notable from this dataset, AAV.CAP-B10 is detargeted from the entire periphery and exhibits almost complete specificity for neurons over other cell types in the CNS. Of the 82,000 variants that comprised our second round of selection, roughly 39,000 exhibited positive enrichment in the brain and negative enrichment in the liver. Among those, the variants tested *in vivo* in mice showed expression patterns that correlated closely with their NGS enrichments and rankings. Furthermore, the presence of additional sequences that were positively enriched in the brain and negatively enriched in peripheral organs indicates the richness of this dataset and engineering location, with many more potentially interesting variants to be identified and characterized. Together, these results indicate that future engineering efforts can also take a stepwise approach toward attaining specificity for certain targets, with each engineering round refining and enhancing novel tropisms identified in the previous round. Importantly, this implies that AAV.CAP-B10 may be further refined in its cell-type specificity to neuronal sub-classes in the CNS.

The therapeutic use of engineered AAVs for treating disease has increased exponentially in recent years, including the use of systemically administered AAVs for CNS gene therapies. The brain is a prime target for gene therapy of a wide array of diseases such as Huntington’s disease, Parkinson’s disease, and Friedreich’s ataxia^49–51^. To achieve the wide coverage and non-invasive delivery needed for effective treatment of these diseases, systemic administration is necessary. Specificity toward the therapeutic target of interest with decreased off-target expression is also necessary to minimize side-effects. For these viruses to find eventual use in gene therapy applications, though, their efficacy needs to be tested and characterized in NHPs. The variants engineered in this study, including AAV.CAP-B10 and AAV.CAP-B22, represent notable progress toward evolving variants that are capable of efficiently, specifically, and safely delivering gene therapies to the CNS in NHPs and, ultimately, humans. The CNS transduction and cell-type bias observed following two rounds of selection in mice were recapitulated in marmosets, indicating that M-CREATE in rodents can be effectively used as a screening platform prior to direct screening or validation in NHPs, greatly increasing the throughput of the engineering process. Together, these results constitute a dramatic step forward toward achieving the goal of engineered AAV vectors that can be used to broadly deliver gene therapies to the CNS in humans.

## Acknowledgements

We thank B. Deverman for helpful discussions on experimental design and implementation. We wish to thank the entire Gradinaru laboratory for helpful discussions. We are grateful to I. Antoshechkin and the Millar and Muriel Jacobs Genetics and Genomics Core at Caltech for assistance with next generation sequencing. We thank M. Smith for helpful discussions and assistance with the bioinformatics pipeline. This work was primarily supported by Defense Advanced Research Projects Agency grant W911NF-17-2-0036 to V.G. and grants from the National Institutes of Health (NIH) to V.G.: Director’s New Innovator DP2NS087949 and PECASE, and BRAIN R01MH117069 and NIH Pioneer DP1OD025535. In addition, this work is funded by the Beckman Institute at Caltech, the Vallee Foundation, and the Moore Foundation. V.G. is a Heritage Principal Investigator supported by the Heritage Medical Research Institute.

## Author information

These authors contributed equally: Nicholas Flytzanis, Nick Goeden

## Affiliations

Division of Biology and Biological Engineering, California Institute of Technology, Pasadena, CA, 91125, USA.

Nicholas Flytzanis, Nick Goeden, David Goertsen & Viviana Gradinaru

National Institute of Mental Health, National Institutes of Health, Bethesda, MD 20892 USA.

Alexander Cummins & James Pickel

## Contributions

N.C.F and N.G. designed and performed experiments and analyzed data. D.G. performed marmoset single variant characterization, analyzed data, and prepared figures. J.P and A.C. performed non-human primate experiments and analyzed the data. J.P. supervised the non-human primate experiments and helped prepare the associated figure. N.C.F., N.G., D.G., and V.G. wrote the manuscript with input from all authors. No experimental work was performed during the Los Angeles COVID-19 stay-at-home order. V.G. supervised all aspects of the work.

## Competing interests

The California Institute of Technology has filed and licensed patent applications for the work described in this manuscript with N.C.F., N.G., and V.G. listed as inventors.

## Corresponding author

Correspondence to

## MATERIALS AND METHODS

### Plasmids

The first-round viral DNA library was generated by amplification of a section of the AAV-PHP.eB capsid genome between amino acids 450 and 599 using NNK degenerate primers (Integrated DNA Technologies, Inc., IDT) to substitute amino acids 452-458 with all possible variations. The resulting library inserts were then introduced into the rAAV-ΔCap-in-cis-Lox plasmid via Gibson assembly as previously described^17^. The resulting capsid DNA library, rAAV-Cap-in-cis-Lox, contained a diversity of ~1.28 billion variants at the amino acid level. The second-round viral DNA library was generated similarly to the first round, but instead of NNK degenerate primers at the 452-458 location, a synthesized oligo pool (Twist Bioscience) was used to generate only selected variants. This second-round DNA library contained a diversity of ~82,000 variants at the amino acid level.

The AAV2/9 REP-AAP-ΔCap plasmid transfected into HEK293T cells for library viral production was modified from the AAV2/9 REP-AAP plasmid previously used^17^ by deletion of the amino acids between 450 and 592. This modification prevents production of a wild-type AAV9 capsid during viral library production after a plausible recombination event between this plasmid co-transfected with rAAV-ΔCap-in-cis-Lox containing the library inserts.

Three rAAV genomes were used in this study. The first, pAAV-CAG-mNeonGreen (Addgene #99134), utilizes a single-stranded (ss) rAAV genome containing the fluorescent protein mNeonGreen under control of the ubiquitous CMV-β-Actin-intron-α-Globin hybrid promoter (CAG). The second, pAAV-CAG-NLS-GFP (Addgene #104061), utilizes an ssAAV genome containing the fluorescent protein EGFP flanked by two nuclear localization sites, PKKKRKV, under control of the CAG promoter. The third, pAAV-CAG-FXN-HA, utilizes an ssAAV genome containing an HA-tagged human frataxin (FXN) protein under control of the CAG promoter and harboring a unique 12bp sequence in the 3’ UTR to differentiate different capsids packaging the same construct.

### Viral production

Recombinant AAVs were generated according to established protocols^52^. Briefly, HEK293T cells (ATCC) were triple transfected using polyethylenimine (PEI); virus was collected after 120 hours from both cell lysates and media and purified over iodixanol (Optiprep, Sigma).

A modified protocol was used for transfection and purification of viral libraries. First, to prevent mosaic capsid formation, only 10 ng of rAAV-Cap-in-cis-Lox library DNA was transfected (per 150-mm plate) to decrease the likelihood of multiple library DNAs entering the same cell. Second, virus was collected after 60 hours, instead of 120 hours, to limit secondary transduction of producer cells. Finally, instead of PEG precipitation of the viral particles from the media, as performed in the standard protocol, media was concentrated >60-fold for loading onto iodixanol.

### Animals

All rodent procedures were approved by the Institutional Animal Care and Use Committee (IACUC) of the California Institute of Technology. Transgenic animals, expressing Cre under the control of various cell-type-specific promoters, and C57Bl/6J WT mice (000664) were purchased from the Jackson Laboratory (JAX). Transgenic mice included Syn1-Cre (3966), GFAP-Cre (012886), and Tek-Cre (8863). For round 1 and round 2 selections from the viral library, we used one male and one female mouse from each transgenic line (aged 8-12 weeks), as well as a single male C57Bl/6J mouse. For validation of individual viral variants, male C57Bl/6J mice aged 6-8 weeks were used. Intravenous administration of rAAV vectors was performed via injection into the retro-orbital sinus.

Marmoset (*Callithrix jacchus*) procedures were approved by ACUC of the National Institutes of Mental Health. Marmosets were born and raised in NIMH colonies and housed in family groups under standard conditions of 27°C temperature and 50% humidity. They were fed ad libitum and received enrichment as part of the primate enrichment program for NHPs at the NIH. For AAV infusions, animals were screened for endogenous neutralizing antibodies (NAb). None of the animals that were screened showed any detectible blocking reaction at 1:5 dilution of serum (Penn Vector Core, University of Pennsylvania). They were then housed individually for several days and acclimated to a new room before injections. Two animals were used for the pooled injection study, both males, aged 7.6 (MPV1) and 11.5 (MPV2) years (Table 1). Five animals were injected with single variants for characterization, but only four were usable (Table 2) as one animal (AAV.CAP-B22 injected, 6.9 years, female, 0.475 kg) was found dead (27 days post-injection); at necropsy the pathology report indicated chronic nephritis unrelated to the virus. The day before infusion the animals’ food was removed. Animals were anesthetized with isoflurane in oxygen, the skin over the femoral vein was shaved and sanitized with an isopropanol scrub, and the virus (Table 1 and Table 2) was infused over several minutes. Anesthesia was withdrawn and the animals were monitored until they became active, upon which they were returned to their cages. Activity and behavior were closely monitored over the next three days, with daily observations thereafter.

### DNA/RNA recovery and sequencing

Round 1 and round 2 viral libraries were injected into C57Bl/6J and Cre-transgenic animals at a dose of 8 × 10^10^ vg/animal and rAAV genomes were recovered two weeks post injection. Mice were euthanized, and most major organs were recovered, snap frozen on dry ice, and placed into long-term storage at −80°C. Tissues collected included: brain, spinal cord, dorsal root ganglia (DRGs), liver, lungs, heart, stomach, intestines, kidneys, spleen, pancreas, testes, skeletal muscle, and adipose tissue. 100 mg of each tissue (~250 mg for brain hemispheres, <100 mg for DRGs) was homogenized in Trizol (Life Technologies, 15596) using a BeadBug (Benchmark Scientific, D1036) and viral DNA was isolated according to the manufacturer’s recommended protocol. Recovered viral DNA was treated with RNase, underwent restriction digestion with SmaI (found within the ITRs) to improve later rAAV genome recovery by PCR, and purified with a Zymo DNA Clean and Concentrator kit (D4033). Viral genomes flipped by Cre-recombinase in select transgenic lines (or pre-flipped in WT animals) were selectively recovered using the following primers: 5’-CTTCCAGTTCAGCTACGAGTTTGAGAAC-3’ and 5’-CAAGTAAAACCTCTACAAATGTGGTAAAATCG-3’, after 25 cycles of 98°C for 10 s, 60°C for 15 s, and 72°C for 40 s, using Q5 DNA polymerase in five 25 μl reactions with 50% of the total extracted viral DNA as a template.

After Zymo DNA purification, samples from the WT C57Bl/6J animals were serially diluted from 1:10 – 1:10,000 and each dilution further amplified around the library variable region. This amplification was done using the primers 5’-ACGCTCTTCCGATCTAATACTTGTACTATCTCTCTAGAACTATT-3’ and 5’-TGTGCTCTTCCGATCTCACACTGAATTTTAGCGTTTG-3’ and 10 cycles of 98°C for 10 s, 61°C for 15 s, and 72°C for 20 s, to recover 73 bp of viral genome around and including the 21 bp variable region and add adapters for Illumina next-generation sequencing. After PCR cleanup, these products were further amplified using NEBNext Dual Index Primers for Illumina sequencing (New England Biolabs, E7600), after 10 cycles of 98°C for 10 s, 60°C for 15 s, and 72°C for 20 s. The amplification products were run on a 2% low-melting-point agarose gel (ThermoFisher Scientific, 16520050) for better separation and recovery of the 210 bp band. The dilution series was analyzed for each WT tissue and the highest concentration dilution which resulted in no product was chosen for further amplification of the viral DNA from the transgenic animal tissues. This process was performed to differentiate between viral genomes flipped prior to packaging or due to Cre in the animal. Pre-flipped viral genomes should be avoided to minimize false positives in the NGS sequencing results.

All Cre-flipped viral genomes from transgenic animal tissues were similarly amplified (using the dilutions that do not produce pre-flipped viral genomes) to add Illumina sequencing adapters and subsequently for index labeling. The amplified products now containing unique indices for each tissue from each animal were run on a low-melting-point agarose gel and the correct bands extracted and purified with a Zymoclean Gel DNA Recovery kit.

Packaged viral library DNA was isolated from the injected viral library by digestion of the viral capsid and purification of the contained ssDNA. These viral genomes were amplified by two PCR amplification steps, like the viral DNA extracted from tissue, to add Illumina adapters and then indices and extracted and purified after gel electrophoresis. This viral library DNA, along with the viral DNA extracted from tissue, was sent for deep sequencing using an Illumina HiSeq 2500 System (Millard and Muriel Jacobs Genetics and Genomics Laboratory, Caltech).

A pool of eight viruses (AAV9, AAV-PHP.eB, AAV.CAP-B1, AAV.CAP-B2, AAV.CAP-B8, AAV.CAP-B10, AAV.CAP-B18, and AAV.CAP-B22) packaging CAG-FXN-HA with unique 12 bp barcodes were injected into two adult marmosets (Table 1). After six weeks, animals were euthanized, and brain and liver were recovered and snap frozen. 1-mm coronal sections from each tissue (4-mm for the brain) were homogenized in Trizol (Life Technologies, 15596) using a BeadBug (Benchmark Scientific, D1036) and total RNA was recovered according to the manufacturer’s recommended protocol. Recovered RNA was treated with DNase, and cDNA was generated from the mRNA using Superscript III (Thermo Fisher Scientific, 18080093) and oligo(dT) primers according to the manufacturer’s recommended protocol. Barcoded FXN transcripts were recovered from the resulting cDNA library, as well as the injected pool, using the primers 5’-TGGACCTAAGCGTTATGACTGGAC-3’ and 5’-GGAGCAACATAGTTAAGAATACCAGTCAATC-3’ and 25 cycles of 98°C for 10 s, 63°C for 15 s, and 72°C for 20 s, using Q5 DNA polymerase in five reactions using 50 ng of cDNA or viral DNA, each, as a template. After Zymo DNA purification, samples were diluted 1:100 and further amplified around the barcode region using the primers 5’-ACGCTCTTCCGATCTTGTTCCAGATTACGCTTGAG-3’ and 5’-TGTGCTCTTCCGATCTTGTAATCCAGAGGTTGATTATCG-3’ and 10 cycles of 98°C for 10 s, 55°C for 15 s, and 72°C for 20 s. After PCR cleanup, these products were further amplified using NEBNext Dual Index Primers for Illumina sequencing (New England Biolabs, E7600), after 10 cycles of 98°C for 10 s, 60°C for 15 s, and 72°C for 20 s. The amplification products were run on a 2% low-melting-point agarose gel (ThermoFisher Scientific, 16520050) for better separation and recovery of the 210 bp band. All indexed samples were sent for deep sequencing as described above.

### NGS data alignment and processing

Raw fastq files from NGS runs were processed with custom-built scripts (codes will be made available on Github) that align the data to an AAV9 template DNA fragment containing the 21 bp diversified region between AA 452-458, for the two rounds of AAV evolution/selection, or to an FXN-HA template containing the 12 bp unique barcode, for the marmoset virus pool. The pipeline to process these datasets involved filtering to remove low-quality reads, utilizing a quality score for each sequence, and eliminating bias from PCR-induced mutations or high GC-content. The filtered dataset was then aligned by a perfect string match algorithm and trimmed to improve the alignment quality. For the AAV engineering, read counts for each sequence were pulled out and displayed along with their enrichment score, defined as the relative abundance of the sequence found within the specific tissue over the relative abundance of that sequence within the injected viral library. For the pooled barcodes, read counts for each sequence were pulled out and normalized to the respective contribution of that barcode to the initial, injected pooled virus to account for small inequalities in the amount of each member of the pool that was injected into the marmosets.

### Tissue preparation and immunohistochemistry

Mice were euthanized with Euthasol and transcardially perfused with ice-cold 1× PBS and then freshly prepared, ice-cold 4% paraformaldehyde (PFA) in 1× PBS. All organs were excised and post-fixed in 4% PFA at 4°C for 48 hours and then sectioned at 50 μm with a vibratome. Immunohistochemistry (IHC) was performed on floating sections with primary and secondary antibodies in PBS containing 10% donkey serum and 0.1% Triton X-100. Primary antibodies used were rabbit anti-NeuN (1:200, Abcam, 177487), rabbit anti-S100 (1:200, Abcam, 868), rabbit anti-Olig2 (1:200, Abcam, 109186), and rabbit anti-Calbindin (1:200, Abcam, 25085). Primary antibody incubations were performed for 16–20 hours at room temperature (RT). The sections were then washed and incubated with secondary Alexa-647 conjugated anti-rabbit FAB fragment antibody (1:200, Jackson ImmunoResearch Laboratories, Inc., 711-607-003) for 6–8 hours at RT. For nuclear staining, floating sections were incubated in PBS containing 0.2% Triton X-100 and DAPI (1:1000, Sigma Aldrich, 10236276001) for 6–8 hours and then washed. Stained sections were then mounted with ProLong Diamond Antifade Mountant (ThermoFisher Scientific, P36970).

Marmosets were euthanized (Euthanasia, VetOne) and perfused with 1× PBS. One hemisphere of the brain (cut into coronal blocks) and the liver were flash frozen in 2-methylbutane (Sigma Aldrich, M32631) chilled with dry ice. The other hemisphere and organs were removed and post-fixed with 4% PFA at 4°C for 48 hours. These organs were then cryoprotected using 10% glycerol followed by 20% glycerol and flash frozen in 2-methylbutane chilled with dry ice. The blocks of tissue were sectioned on an AO sliding microtome. 50-μm sections were collected in PBS and IHC was performed on floating sections. To visualize transduced cells expressing the HA-tagged FXN from the variant pool, sections were incubated overnight at RT with a rabbit anti-HA primary antibody (Cell Signaling Technologies, C29F4). Following primary incubation, sections were washed in PBS and then incubated with a biotinylated goat anti-rabbit secondary antibody (1:200, Vector Laboratories, BA1000) for one hour at RT. Sections were again washed in PBS and incubated for 2 hours in ABC Elite (Vector Laboratories, PK6100) as outlined by the supplier. The ABC Peroxidase complex was visualized using 3,3’-Diaminobenzidine tetrahydrochloride hydrate (DAB, Sigma Aldrich, D5637) for 5 minutes at RT. Sections were then mounted for visualization.

To visualize transduced cells expressing the HA-tagged FXN for individual variant injections, immunohistochemistry staining was performed on floating sections with primary and secondary antibodies in PBS containing 10% donkey serum and 0.1% Triton X-100. Primary antibodies used were rat anti-HA (1:200, Roche, 3F10), rabbit anti-NeuN (1:200, Abcam, 177487), and rabbit anti-S100 beta (1:200, Abcam, 52642). Primary antibody incubations were performed for 16–20 hours at RT. The sections were then washed and incubated with secondary anti-rat Alexa-488 (1:200, ThermoFisher, A-21208) and anti-rabbit Alexa-647 (1:200, ThermoFisher, A32795). Stained sections were then washed with PBS and mounted with ProLong Diamond Antifade Mountant.

### Imaging and quantification

All CAG-mNeonGreen-expressing tissues were imaged on a Zeiss LSM 880 confocal microscope using a Fluar 5× 0.25 M27 objective, with matched laser powers, gains, and gamma across all samples of the same tissue. The acquired images were processed in Zen Black 2.3 SP1 (Zeiss).

All CAG-NLS-GFP-expressing tissues were imaged on a Keyence BZ-X all-in-one fluorescence microscope at 48-bit resolution with the following objectives: PlanApo-λ 20x/0/75 (1-mm working distance) or PlanApo-λ 10x/0.45 (4-mm working distance). For colocalization of GFP expression to antibody staining, in some cases the exposure time for the green (GFP) channel was adjusted to facilitate imaging of high- and low-expressing cells while avoiding oversaturation. In all cases in which fluorescence intensity was compared between samples, exposure settings and changes to gamma or contrast were maintained across images. To minimize bias, multiple fields of view per brain region and peripheral organ were acquired for each sample. For brain regions, the fields of view were matched between samples and chosen on the basis of the antibody staining rather than GFP signal. For peripheral tissues, fields of view were chosen on the basis of the DAPI or antibody staining to preclude observer bias.

Marmoset tissue sections transduced with the variant pool were examined and imaged on a Zeiss AxioImager Z1 with an Axiocam 506 color camera. Acquired images were processed in Zen Blue 2 (Zeiss).

Marmoset tissues transduced with individual variants were imaged on a Zeiss LSM 880 confocal microscope. Whole tissue sections were imaged using a Fluar 5× 0.25 M27 objective. Images for cortex quantification were imaged with a LD-LCI Plan-Apochromat 25x/0.8 M27 objective. Imaging at each magnification was performed with matched laser powers, gains, and gamma across all samples of the same tissue. The acquired images were processed in Zen Black 2.3 SP1 (Zeiss). Quantification of transduction in marmoset cortex was performed through manual cell counting.

All image processing was performed with the Keyence BZ-X Analyzer. Colocalization between the GFP signal and antibody or DAPI staining was performed using the Keyence BZ-X Analyzer with the hybrid cell count automated plugin. Automated counts were validated and routinely monitored by comparison with manual hand counts and found to be below the margin of error for manual counts.

To compare total cell counts and fluorescence intensity throughout the brain between samples, an entire sagittal section located 1200 μm from the midline was imaged using matched exposure conditions with the Keyence BZ-X automated XY stitching module. Stitched images were then deconstructed in the Keyence BZ-X Analyzer suite and run through the hybrid cell count automated plugin to count the total number of cells in the entire sagittal section. Average fluorescence intensity was calculated by creating a mask of all GFP-positive cells throughout the sagittal section and measuring the integrated pixel intensity of that mask. The total integrated pixel intensity was divided by the total cell count to obtain the fluorescence intensity per cell. In all cases where direct comparisons of fluorescence intensity were made, exposure settings and post-processing contrast adjustments were matched between samples.

### Statistics

Microsoft Excel 2018 and GraphPad Prism 8 were used for statistical analysis and data representation. Unless otherwise noted, all experimental groups were n = 6, determined using preliminary data and experimental power analysis. For the statistical analyses and related graphs, a single data point was defined as two tissue sections per animal, with multiple technical replicates per section when possible. Technical replicates were defined as multiple fields of view per section, with the following numbers for each region or tissue of interest: cerebellum = 3, cortex = 4, hippocampus = 3, midbrain = 1, striatum = 3, thalamus = 4, liver = 4, spleen = 2, testis = 2, kidney = 2, lung = 2, spine = 1, DRG = 1, whole sagittal = 1.

For all statistical analyses, significance is represented as: *P ≤ 0.05; **P ≤ 0.01; ***P ≤ 0.001; ****P ≤ 0.0001; n.s., P ≥ 0.05.

### Data availability

All constructs and tools will be available through Addgene and the Beckman Institute CLOVER center: clover.caltech.edu. The data that support the findings of this study are available from the corresponding author upon request.

