## Supplementary Figures for "Broad gene expression throughout the mouse and marmoset brain after intravenous delivery of engineered AAV capsids"

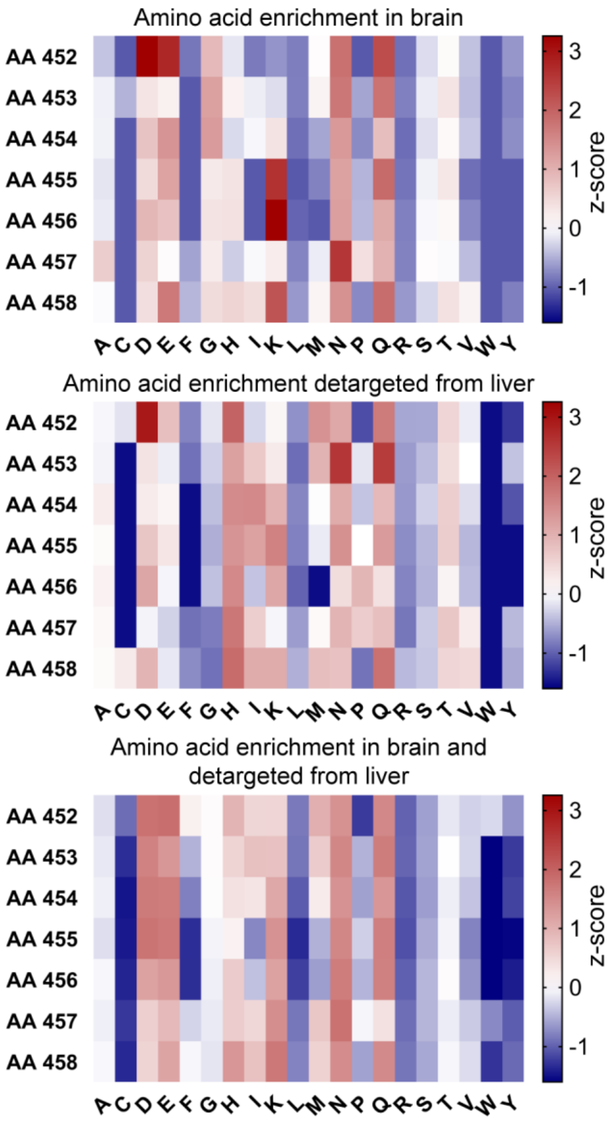


**Figure S1. Related to Figure 1.** Amino-acid contributions across the substitution library.
Amino-acid contributions across the 7-mer substitution to variants enriched in the brain and detargeted from the liver. The 1000 variants with the highest enrichment in the brain of hSyn-Cre animals, the 1000 variants with the lowest enrichment in the liver of Tek-Cre animals, and all variants with positive enrichment in the brain and negative enrichment in the liver were analyzed. Plotted is the z-score of all amino acids at each position. Amino acids found at a higher prevalence than average are displayed in red and those found at a lower prevalence are displayed in blue.

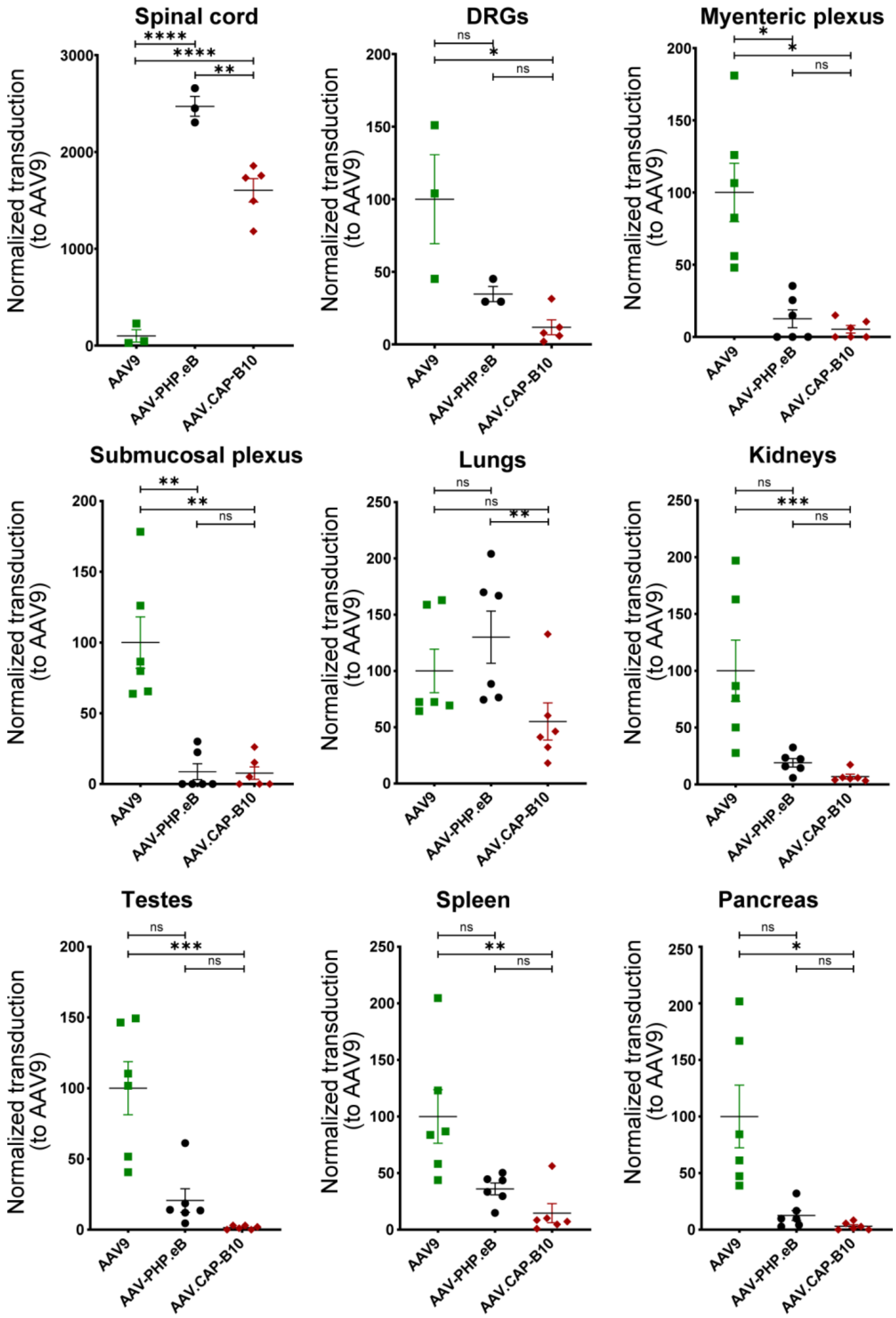


**Figure S2. Related to Figure 2.** Quantification of detargeting from peripheral organs in mice. ssAAV9:CAG-NLSx2-EGFP, ssAAV-PHP.eB:CAG-NLSx2-EGFP, and ssAAV.CAP-B10:CAG-NLSx2-EGFP were intravenously injected into male adult mice at 1 × $10^{11}$ vg/mouse. GFP fluorescence was assessed after three weeks of expression. Transduction efficiency of AAV.CAP-B10 in the spinal cord is significantly increased relative to AAV9 and significantly decreased relative to AAV-PHP.eB. Transduction efficiency of AAV.CAP-B10 in the DRGs is significantly decreased relative to AAV9 and non-significantly decreased relative to AAV-PHP.eB. In the myenteric and submucosal plexi of the intestines, AAV.CAP-B10 transduction efficiency is significantly decreased relative to AAV9 and non-significantly decreased relative to AAV-PHP.eB. In the lungs, AAV.CAP-B10 transduction efficiency is significantly decreased relative to AAV-PHP.eB and non-significantly decreased relative to AAV9. In the kidneys, spleen, pancreas, and testes, AAV.CAP-B10 transduction efficiency is significantly decreased relative to AAV9 and non-significantly decreased relative to AAV-PHP.eB. n = 6 mice per group except for the spinal cord and DRGs, which had n = 3 mice for the AAV9 and AAV-PHP.eB groups and n

= 5 mice for the AAV.CAP-B10 group; mean ± SE; ANOVA for spinal cord, Brown-Forsythe and Welch ANOVA tests for myenteric plexus and pancreas, Kruskal-Wallis test for DRGs, submucosal plexus, lungs, kidneys, spleen, and testes (*P ≤ 0.05; **P ≤ 0.01; ***P ≤ 0.001; ****P ≤ 0.0001; n.s., P ≥ 0.05).

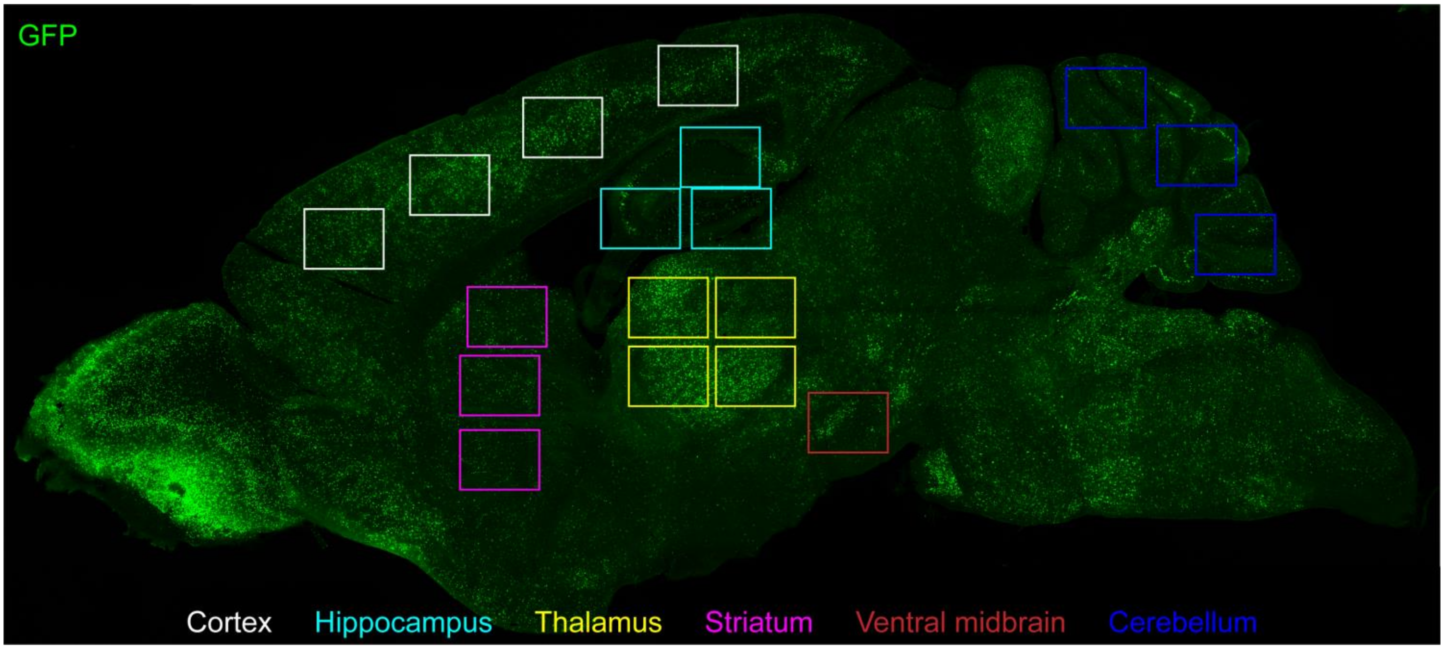


**Figure S3. Related to Figure 2.** Brain regions selected for cell-type quantification. Four areas of cortex (white), three areas of hippocampus (teal), four areas of thalamus (yellow), three areas of striatum (purple), one area of ventral midbrain (red), and three areas of cerebellum (dark blue) were selected from each brain section quantified. Each section was located roughly 1200 µm from the midline for consistency across animals.

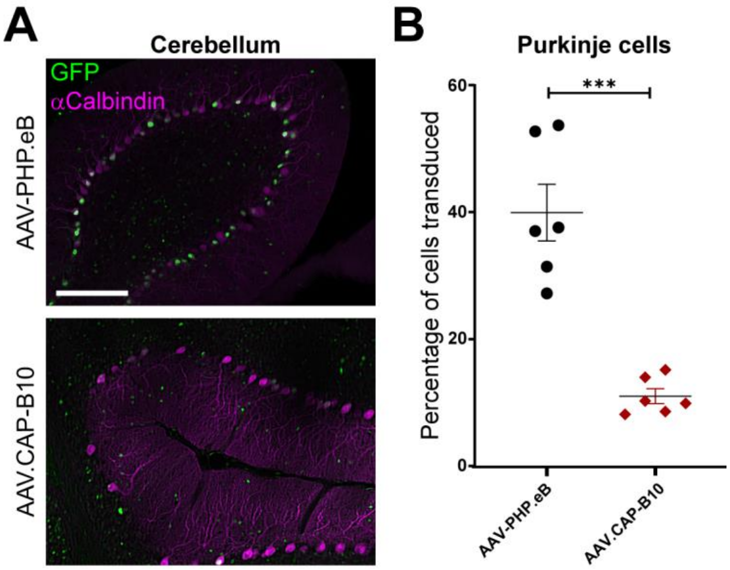


**Figure S4. Related to Figure 2.** Quantification of Purkinje cell expression. AAV.CAP-B10 is significantly detargeted from Purkinje cells in the cerebellum. ssAAV-PHP.eB:CAG-NLSx2-EGFP and ssAAV.CAP-B10:CAG-NLSx2-EGFP were intravenously injected into male adult mice at $1 \times 10^{11}$ vg/mouse. GFP fluorescence was assessed after three weeks of expression. (**a, b**) Quantification of Purkinje cell transduction in the cerebellum shows significantly fewer cells transduced by AAV.CAP-B10 than by AAV-PHP.eB. n = 6 mice per group, mean ± SE, Welch's t test (***P ≤ 0.001). Scale bar is 200 µm.

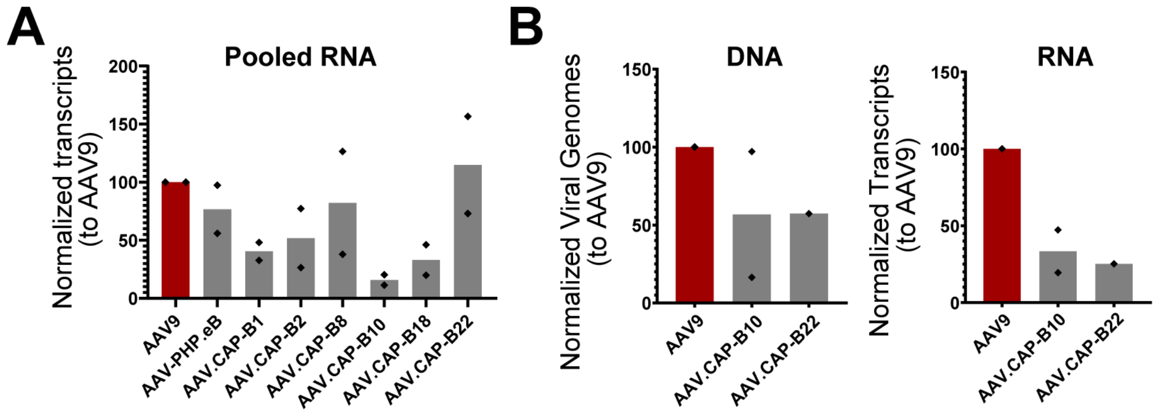


**Figure S5. Related to Figure 3 and Figure 4.** Marmoset liver profiling. (**a**) NGS quantification of pooled variant injections, showing relative detargeting from the liver for the different variants. AAV.CAP-B22 contributes similar RNA levels as AAV9, but AAV.CAP-B10 contributes >5-fold less. n = 2 marmosets. (**b**) Viral genome and RNA transcript measurements in liver tissue after single variant injection. Results indicate that AAV.CAP-B10 and AAV.CAP-B22 have lower transduction in the liver than AAV9. Total values for viral genomes were normalized by tissue mass, whereas RNA transcripts were normalized by the total number of transcripts.

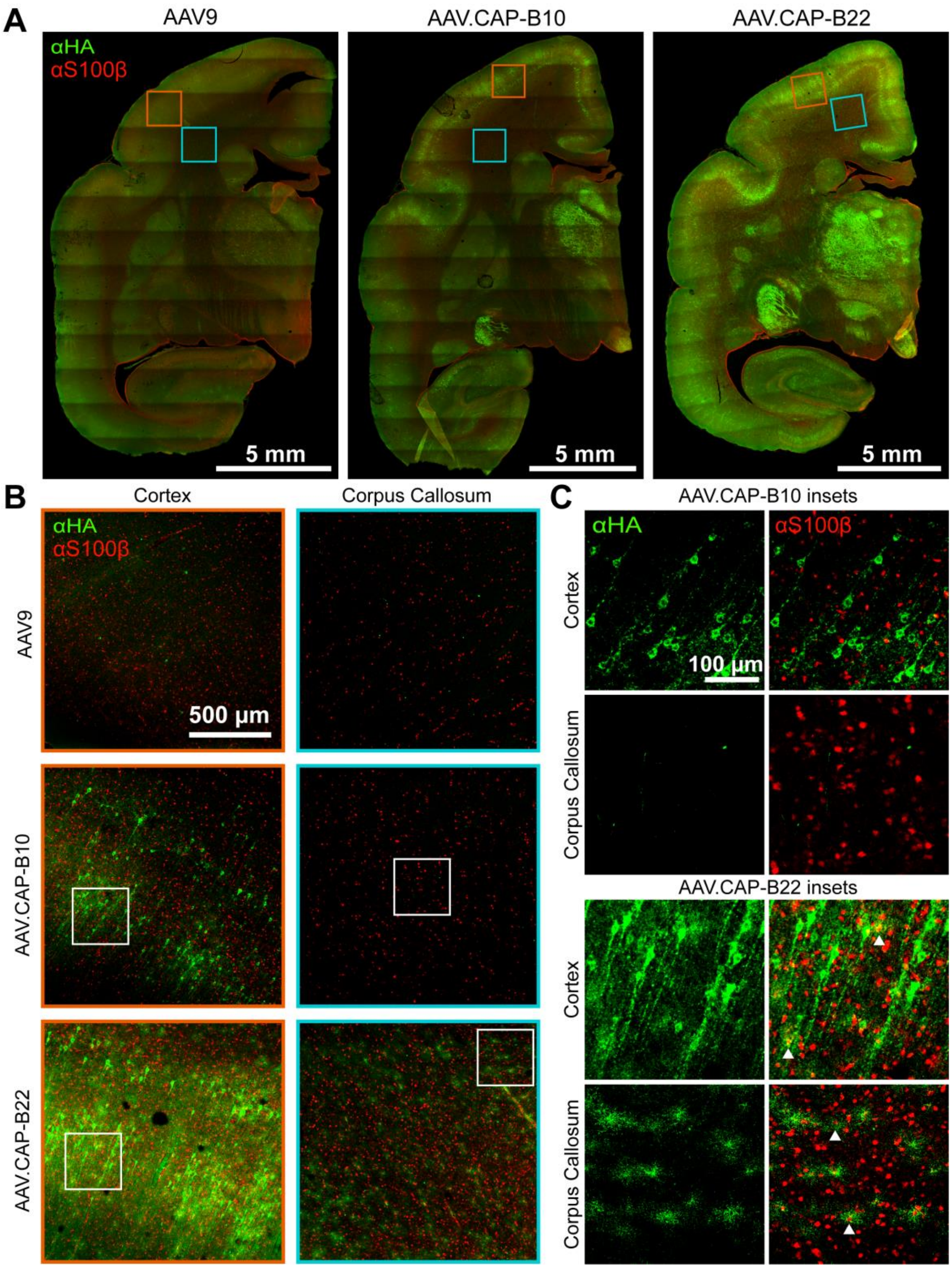


**Figure S6. Related to Figure 4.** Astrocyte specificity of AAV9, AAV.CAP-B10, and AAV.CAP-B22 was characterized through immunostaining for the HA tag in conjunction with S100β. (**a**) Coronal sections from marmoset brains after injection of individual variants, showing the locations of the areas selected for magnification. (**b**) Transduction of astrocytes was compared in magnified frames taken from the cortex and corpus callosum of animals injected with single variants. AAV9 and AAV.CAP-B10 display very little astrocyte transduction, whereas AAV.CAP-B22 displays transduction of astrocytes in both the cortex and the corpus callosum. (**c**) Further magnification indicates colocalization (examples marked with arrows) of HA and S100β immunostaining in both the cortex and corpus callosum of marmosets for AAV.CAP-B10 and AAV.CAP-B22.
